# Cellular deconvolution of GTEx tissues powers eQTL studies to discover thousands of novel disease and cell-type associated regulatory variants

**DOI:** 10.1101/671040

**Authors:** Margaret K. R. Donovan, Agnieszka D’Antonio-Chronowska, Matteo D’Antonio, Kelly A. Frazer

## Abstract

The Genotype-Tissue Expression (GTEx) resource has contributed a wealth of novel insights into the regulatory impact of genetic variation on gene expression across human tissues, however thus far has not been utilized to study how variation acts at the resolution of the different cell types composing the tissues. To address this gap, using liver and skin as a proof-of-concept tissues, we show that readily available signature genes based on expression profiles of mouse cell types can be used to deconvolute the cellular composition of human GTEx tissues. We then deconvoluted 6,829 bulk RNA-seq samples corresponding to 28 GTEx tissues and show that we are able to quantify cellular heterogeneity, determining both the different cell types present in each of the tissues and how their proportions vary between samples of the same tissue type. Conducting eQTL analyses for GTEx liver and skin samples using cell type composition estimates as interaction terms, we identified thousands of novel genetic associations that had lower effect sizes and were cell-type-associated. We further show that cell-type-associated eQTLs in skin colocalize with melanoma, malignant neoplasm, and infection signatures, indicating variants that influence gene expression in distinct skin cell types play important roles in skin traits and disease. Overall, our study provides a framework to estimate the relative fractions of different cell types in GTEx tissues using signature genes from mouse cell types and functionally characterize human genetic variation that impacts gene expression in a cell-type-specific manner.

## Introduction

Understanding the regulatory impact of genetic variation on complex traits and disease has been a longstanding goal of the field of human genetics. To decipher the mechanistic underpinnings of complex traits, the GTEx Project^1^ has generated a large dataset, including over 10,000 bulk RNA-seq samples representing 53 different tissues (corresponding to 30 organs) obtained from 635 genotyped individuals, to link the influence of genetic variants on gene expression levels through expression quantitative trait loci analysis (eQTL). While GTEx has provided important biological insights, unaccounted for cellular heterogeneity (i.e., different cell types within a tissue and the relative proportions of each cell type across samples of the same tissue) present in bulk RNA-seq can affect genotype-gene expression associations^2^. Since regulation of gene expression varies across cell types, not accounting for cellular composition could result in loss or distortion of signal from relatively rare cell types, thus characterization of cellular heterogeneity across all GTEx tissues is critical for more comprehensive eQTL studies. It is possible that future studies pursuing cell-type-associated eQTLs may utilize single cell approaches (e.g. single cell RNA-seq; scRNA-seq); however, non-trivial technical challenges, such as hard to dissociate tissues and low capture efficiencies, make the generation of a GTEx-scale single-cell expression dataset a substantial undertaking, which would take years to complete. Thus, as single-cell large-scale scRNA-seq collections progress, our present knowledge of how genetic variation influences cell-type-associated gene expression would greatly benefit from conducting eQTL analyses on bulk GTEx tissue samples whose cellular heterogeneity has been characterized through existing deconvolution methods^3–5^.

To characterize the heterogeneity of bulk RNA-seq samples, gene signatures from cell types known to be present in a given tissue can be used to deconvolute the cellular composition (i.e. the proportion of each cell type). The signature genes needed to deconvolute a heterogeneous tissue can be obtained by analyzing scRNA-seq generated from an analogous tissue. However, there are relatively few human scRNA-seq resources currently available^6–10^, and thus only a small fraction of GTEx tissues could be deconvoluted using gene expression signatures derived from existing human single-cell data. While human single-cell data is limited, the Tabula Muris exists^11^, which is a powerful resource of scRNA-seq data from mouse including more than 100,000 cells from 20 tissue types (referred in the Tabula Muris resource as organs and tissues). A recent study showed that similar cell types in humans and mice share sufficient gene expression signatures to integrate scRNA-seq data between the two species^12^, raising the possibility of utilizing the available scRNA-seq from mouse to generate the gene expression signatures for deconvolution of GTEx tissues.

To examine the feasibility of using mouse-derived gene expression signatures to deconvolute human tissues, we compared cellular composition estimates of GTEx liver and GTEx skin samples generated using human scRNA-seq to those generated using the Tabula Muris scRNA-seq resource. We show that the human and mouse single-cell data captured many overlapping cell populations and that using either human-derived or mouse-derived gene signatures to deconvolute the 175 GTEx liver samples and the 860 GTEx skin samples resulted in highly correlated estimated cellular compositions. We show that the main differences between the cell types identified using the human-derived versus mouse-derived signature genes were due to: 1) subtle biological differences that exist in human and mouse immune cells, and 2) resolution (i.e., the ability to detect less abundant cell types and distinguish between similar cell types) which was impacted by technical differences in the human and mouse scRNA-seq data sets, including the number of cells captured and subjected to scRNA-seq and the spatial location from which the tissue was sampled. We used gene signatures derived from the Tabula Muris resource to deconvolute 6,829 GTEx samples corresponding to 28 tissues from 14 organs, which enabled us to determine how the fractions of different cell types vary across GTEx samples derived from the same tissue. Using deconvoluted liver and skin GTEx samples for eQTL analyses, we identified thousands of novel (i.e. not detected using bulk RNA-seq samples) genetic associations that tended to have lower effect sizes, some of which are cell-type-associated. Finally, we show that skin cell-type-associated eQTLs colocalize with GWAS variants for melanoma, malignant neoplasm, and infection signatures, indicating that variants that are functional in limited skin cell types may play major roles in skin traits and disease. Taken together, our study demonstrates two major principles: 1) mouse-derived signature genes can be used to deconvolute the cellular composition of human tissues; and 2) the estimation of cellular heterogeneity by deconvolution enhances the genetic insights yielded from the GTEx resource.

## Results

### scRNA-seq from mouse and human analogous tissues capture similar cell types

To examine the extent to which scRNA-seq generated from analogous human and mouse tissues (Table S1) captured similar cell types, we first examined liver as a proof-of-concept tissue (Figure 1A, “proof-of-concept”). We used previously defined cell types from Tabula Muris mouse liver cells (which were purified for viable hepatocyte and non-parenchymal cells followed by FACS sorting; 710 cells; 5 cell types)^11^, and to be consistent, we used the Tabula Muris annotation approach to analyze existing human liver scRNA-seq data (total liver homogenate; 8,119 cells; 15 cell types)^6^. In brief, on the 8,119 human liver single-cells, we performed nearest-neighbor graph-based clustering on components computed from principal component analysis (PCA) of variably expressed genes, and then used marker genes to define the cell populations corresponding to each of the 15 previously observed cell types ^6^ (see Methods). Human and mouse scRNA-seq from liver captured several shared cell types, including hepatocytes, endothelial cells, and various immune cells (Kuppfer cells, B cells, and natural killer (NK) cells) (Figure 1B-E), however we noted that there were many more distinct cell types for human liver. This was due to the fact that cell type resolution (i.e. the ability to distinguish between similar cell types) can be influenced by 1): the number of cells captured and subjected to scRNA-seq, which may influence the proportion of observed common or rare cell types^13^; and 2) how the tissue was sampled, which may enrich for selected populations or capture how populations are distinguished by spatial location (i.e. zonation). Some of the 15 cell types identified in the human liver scRNA-seq were highly similar and clustered near each other, for example there were four hepatocytes populations and two endothelial cell populations (human periportal sinusoidal endothelial cells (SEC) and central venous SECs) distinguished by their zonation (Figure 1B,C). In contrast, for the mouse liver scRNA-seq, which had considerably fewer cells analyzed, we only observed one hepatocyte population and one endothelial population (Figure 1D,E). If we collapsed the cell types that were similar to each other in the human scRNA-seq, we obtained 7 distinct cell classes (Figure 1B,F; Table S2), which largely corresponded to the 5 cell types from mouse liver scRNA-seq (cholangiocytes and hepatic stellate cells were absent due to having been sorted by FACS; Figure D-F). Overall, these results show that scRNA-seq generated from human and mouse liver captured similar cell types and that technical differences, including the number of cells analyzed and tissue sampling methodology, affects the cell type resolution.

**Figure 1:**
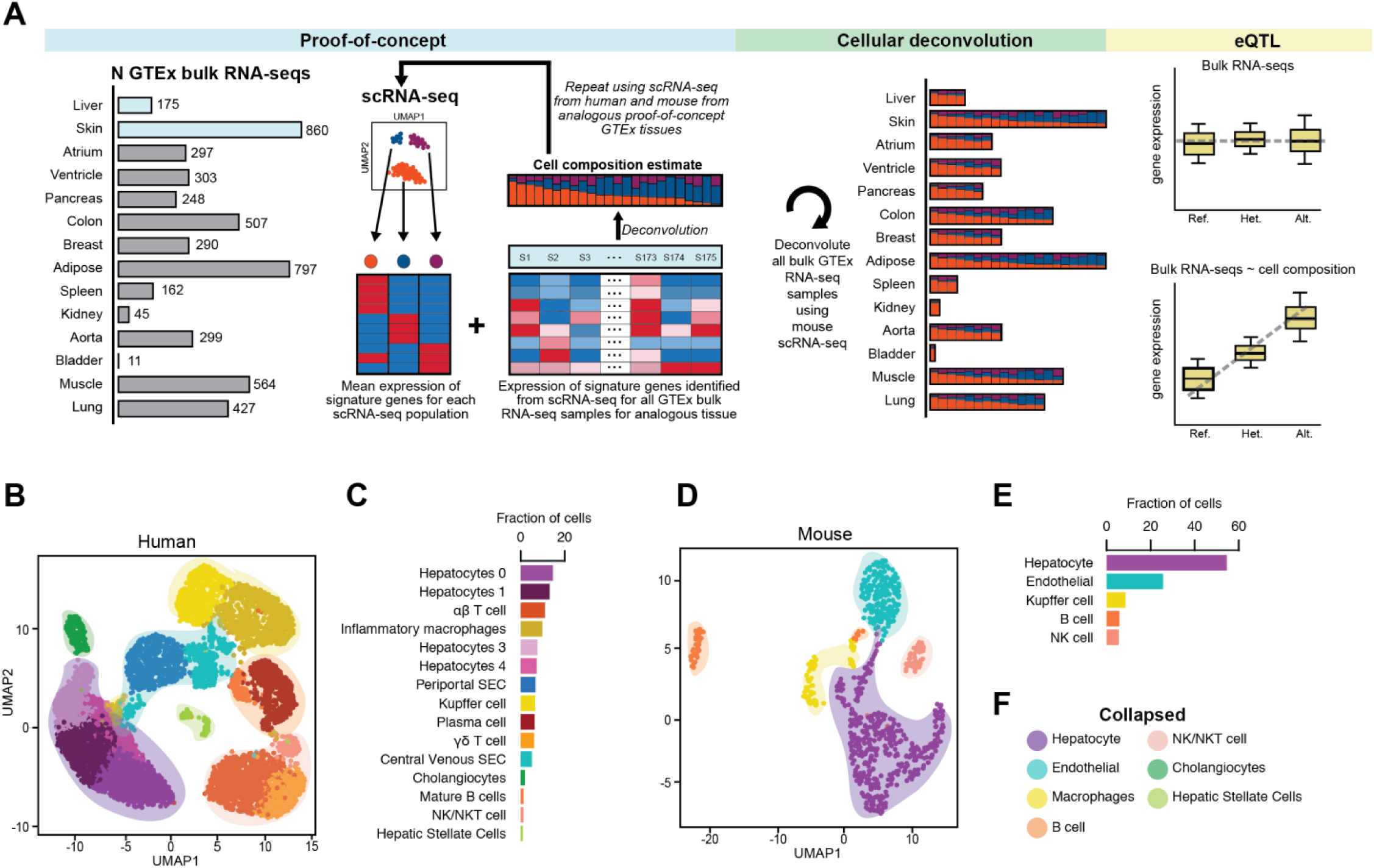
Human and mouse liver scRNA-seq contains similar cell types. A. Overview of the study design. Our goal was to deconvolute the cellular composition of 28 GTEx tissues from 14 organs using mouse scRNA-seq for the purpose of identifying cell-type-associated eQTLs. We first conducted proof-of-concept analyses, where we compared cellular estimates of two proof-of-concept GTEx tissues (liver and skin) after having deconvoluted each using both mouse and human signature genes obtained from scRNA-seq. We then performed cellular deconvolution of the 28 GTEx tissues from 14 organs using CIBERSORT and characterized both the heterogeneity in cellular composition between tissues and the heterogeneity in relative distributions of cell populations between RNA-seq samples from a given tissue. Finally, we used the cell type composition estimates as interaction terms for eQTL analyses to determine if we could detect novel cell-type-associated genetic associations. B. UMAP plot of clustered scRNA-seq data from human liver. Each point represents a single cell and color coding of cell type populations (See Methods: Defining the cellular composition of liver) are shown adjacent (Figure 1C). Similar cell types can be collapsed to single cell type classifications and are noted with colored, transparent shading (Figure 1F). C. Bar plots showing the fraction of each cell type from the scRNA-seq data from human liver. Color-coding of cell types correspond to the colors of the single cells in Figure 1B. D. UMAP plot of clustered scRNA-seq data from mouse liver. Each point represents a single cell and color coding of cell type populations are shown adjacent (Figure 1E). Each cell type has a corresponding collapsed cell type in human liver and is noted with colored, transparent shading (Figure 1F). E. Bar plots showing the fraction of each cell type from the scRNA-seq data from mouse liver. Color-coding of cell types correspond to the colors of the single cells in Figure 1D. F. Legend showing the colors of collapsed similar cell types from human liver (transparent shading in UMAP Figures 1B,D; Table S2). All cell types from mouse liver have a corresponding collapsed cell type in human liver (hepatocyte, endothelial, macrophages, B cell, NK/NKT cell) and human liver also contains two additional cell types not present in mouse (cholangiocytes and hepatic stellate cells).

### Mouse liver signature genes can estimate cellular composition of human liver samples

To establish the ability to use expression profiles of signature genes derived from mouse scRNA-seq for the deconvolution of human GTEx tissues, we first examined if the similarly annotated cell types identified in the two species (Figure 1B-1E) clustered together based on their gene expression profiles. We harmonized the human and mouse liver scRNA-seq using canonical correlation analysis (CCA) and visualized using uniform manifold approximation and projection (UMAP) (Figure 2A,B). We observed that the corresponding cell types across the two species clustered closely together, indicating that they had highly similar gene expression profiles.

**Figure 2:**
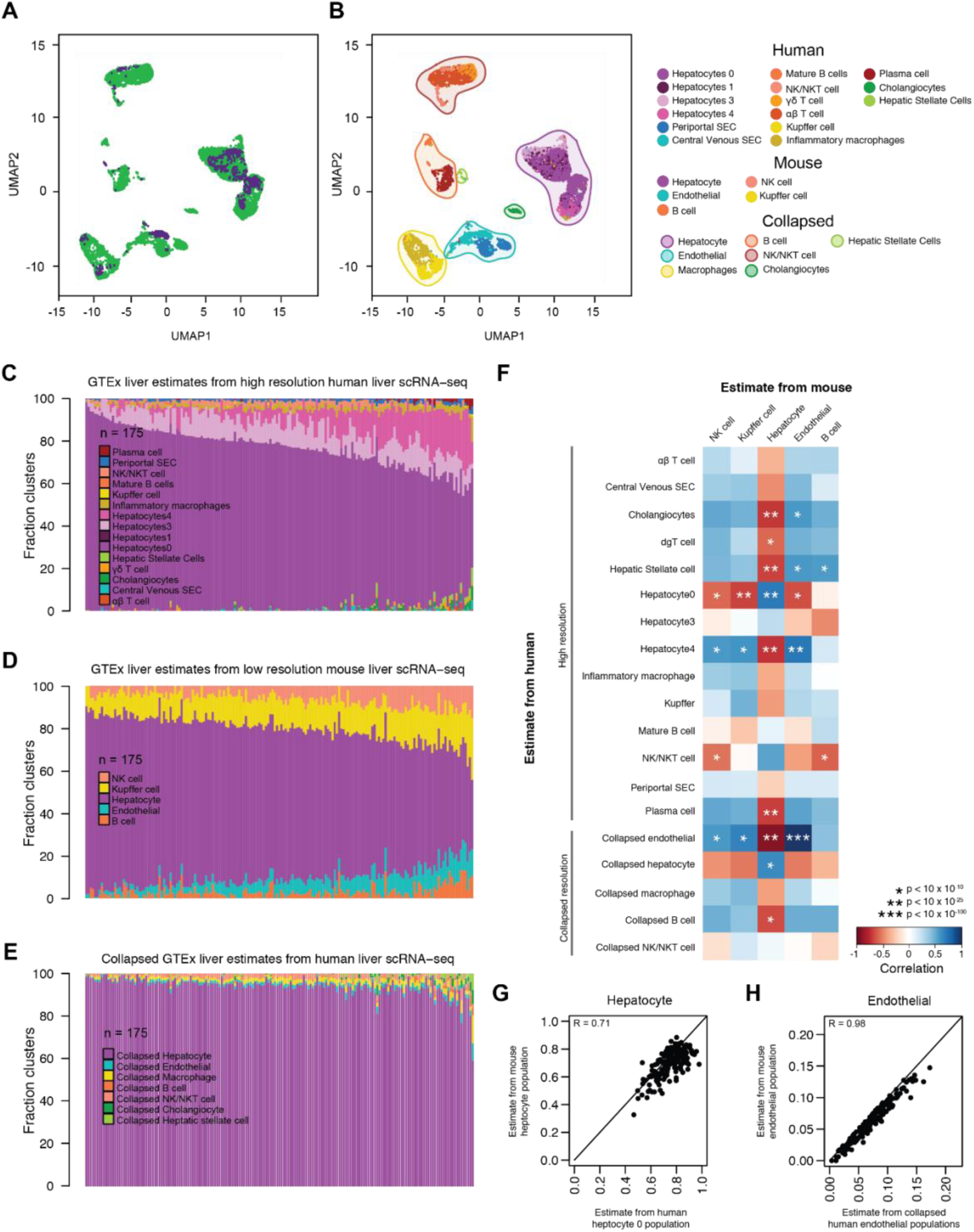
Comparison of GTEx liver cell estimates using mouse versus human signature gene. A. UMAP plot of integrated scRNA-seq data from human and mouse liver. Each point represents a single cell and color coding of cells indicates the species the cells were obtained from (human = green; mouse = purple). B. UMAP plot of integrated scRNA-seq data from human and mouse liver. Each point represents a single cell and color coding of cell type populations are shown in the adjacent legend. The collapsed populations are the same as those shown in Figure 1F. C, D, E. Bar plots showing the fraction of cell types estimated in the 175 GTEx liver RNA-seq samples deconvoluted using gene expression profiles from high resolution human liver scRNA-seq (C), low resolution mouse liver scRNA-seq (D), and GTEx estimates generated by collapsing high resolution human cell types within each of the 7 distinct cell classes (E). F. Heatmap showing the correlation of GTEx liver cell population estimates from human liver scRNA-seq at high and collapsed resolutions (rows) and mouse liver (columns) at low resolution. Color coding of heatmap scales from red, indicating negative correlation in estimates, to blue, indicating positive correlation in estimates. Significance is indicated with asterisks. G, H. Scatter plots of estimated cell compositions across 175 GTEx livers deconvoluted using human scRNA-seq for human hepatocyte 0 population (d) and human collapsed endothelial cells (e) versus estimated cell populations deconvoluted using mouse scRNA-seq.

We next compared the cellular composition estimates of 175 GTEx bulk liver RNA-seq samples^1^ obtained by deconvolution using human signature genes to those obtained using mouse signature genes (Figure 1A, “proof-of-concept”), which respectively consisted of the top 200 most significantly overexpressed genes for each cell type identified in scRNA-seq from high resolution human liver (i.e. signature genes from 15 cell types) and low resolution mouse liver (i.e. signature genes from 5 cell types) (Table S3). From the 175 GTEx bulk liver RNA-seq samples, we independently extracted the expression of the signature genes at the two resolutions, and used CIBERSORT^3^ to estimate the cellular compositions (i.e. high resolution human liver estimates and low resolution mouse liver estimates) (Figure 2C,D; Table S12,13). To investigate how resolution impacted the correlation between human and mouse signature gene estimates, we also collapsed the high resolution human liver cellular composition estimates for each of the 175 deconvoluted samples by summing the estimates across similar cell types in each of the 7 distinct cell classes (Table S2) (Figures 1B,F and 2E). We then calculated all pairwise-correlations between each of the estimated cell populations in the 175 GTEx liver samples from human (high and collapsed resolution estimates) with the estimated cell populations from mouse (low resolution estimates) (Figure 2F, Figure S1). We found that hepatocyte estimates from mouse liver were positively and highly correlated with the human high resolution hepatocyte 0 population estimate (r = 0.71, p-value = 5.4×10^-28^), but not correlated with any of the other three high resolution hepatocyte populations (1, 3 and 4); and was slightly less correlated with the collapsed hepatocyte population estimate (r=0.64, p-value = 1.015×10^-21^) (Figure 2F,G). This indicates that the low resolution mouse hepatocyte population corresponds to one of the four human hepatocyte populations/zones potentially due to tissue sampling. Further, we observed that the endothelial estimates from mouse were highly correlated with the collapsed human endothelial population estimates (r = 0.98, p-value = 1.2×10^-115^) but not correlated with either high resolution human periportal SECs or central venous SECs (Figure 2H). This indicates that the human endothelial population estimates captured a higher resolution of cell type specificity (i.e. two independent endothelial zones), whereas the mouse endothelial population estimates likely captured a mixture of both cell types (i.e. the two endothelial zones are combined into a single cell population), which is potentially due to the lower number of mouse cells analyzed.

While in general we observed high correlation in the human and mouse population estimates for most cell types (hepatocytes, endothelial cells, and Kupffer cells), B cells were non-significantly correlated, and NK-like cells were negatively correlated (Figure 2F). This difference in immune cell estimates in GTEx liver is not wholly unexpected, as biological differences, including immune response differences, exist between species ^14^. Our results show that, while technical differences in scRNA-seq generation and biological differences between humans and mice may impact cell estimation performance, overall mouse signature genes can be used to deconvolute human GTEx bulk RNA-seq samples.

### Deconvolution of GTEx skin confirms mouse signature genes can estimate cellular composition

To examine the similarity of cellular estimates across 860 GTEx human skin samples obtained using human-derived versus mouse-derived signature genes, we used scRNA-seq from human epidermal cells^15^ (digested dorsal forearm skin biopsies; 5,670 cells; 9 cell types) (Figure 3A,B) and Tabula Muris mouse skin cells (FACS sorted epidermal keratinocytes; 2,263 cells; 6 cell types) (Figure 3C,D). While the previous human and mouse liver scRNA-seq studies^6,11^ used similar naming conventions for the cell type annotations (Figure 1B-E), the human and mouse skin scRNA-seq studies^11, 15^ did not (Figure 3A-D), and thus we first needed to identify the corresponding cell types across the two species. To accomplish this, we harmonized the human dermis and mouse skin scRNA-seq using CCA (Figure 3E,F) and visualized using UMAP. We observed three distinct superpopulations: 1) superpopulation 1, epidermal cells, consisting of the four human keratinocyte populations (14, 5, 711, and 1) and mouse epidermal cells, basal cells, stem cells of epidermis, and outer bulge cells (keratinocyte stem cells), 2) superpopulation 2, consisting of the three human fibroblast populations (0, 3, and and mouse inner bulge cells (keratinocyte stem cells); and 3) superpopulation 3, leukocytes, consisting of human lymphocytes and mouse leukocytes (Figure 3E,F). Further, the different cell types within each of the three clusters expressed corresponding marker genes (Figure S2C,F), confirming that they indeed were similar cell types in the human and mouse skin scRNA-seq. Overall, we found human and mouse skin scRNA-seq captured shared cell types that cluster into three distinct superpopulations.

**Figure 3:**
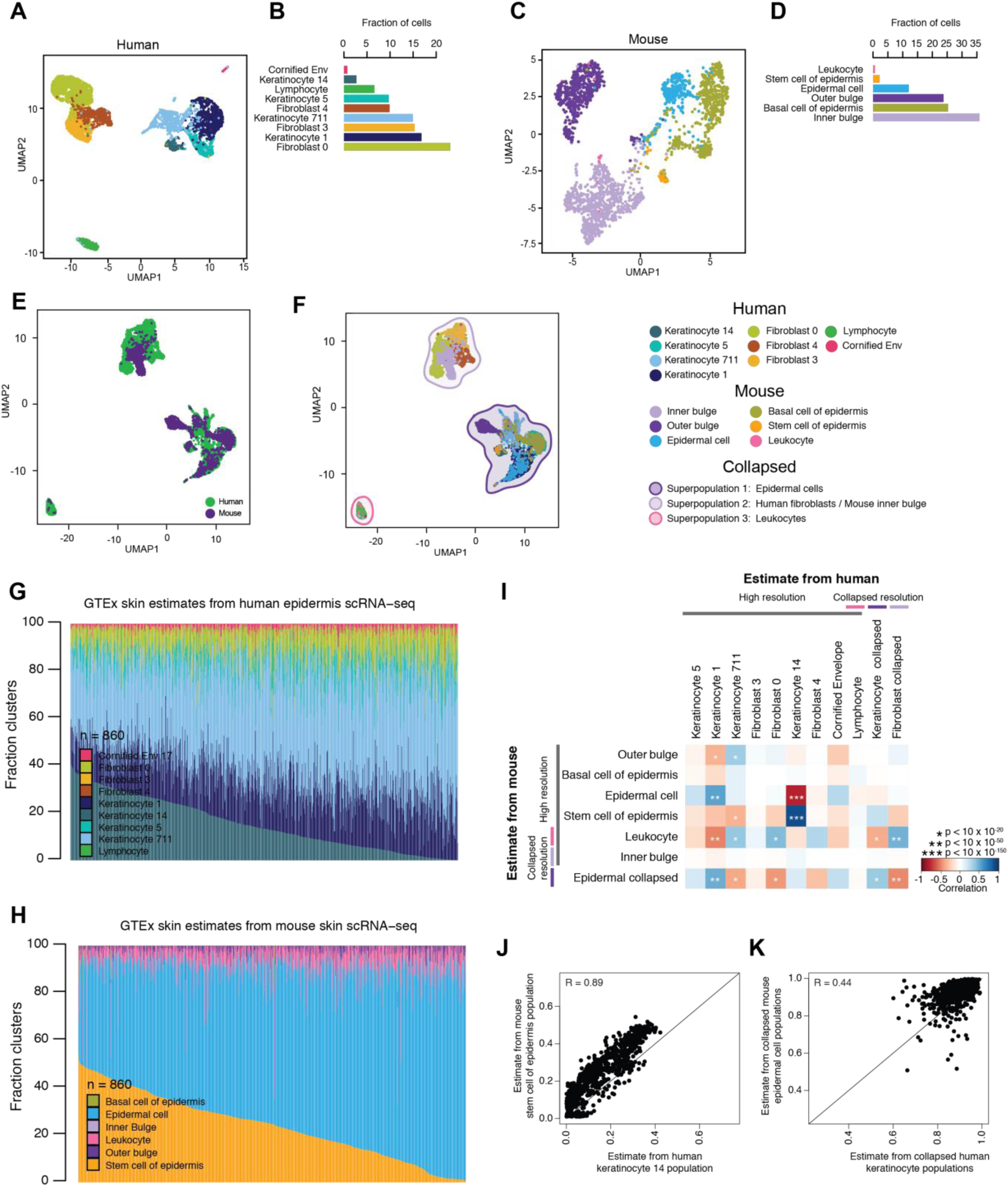
Comparison of GTEx skin cell estimates using mouse versus human signature genes. A. UMAP plot of clustered scRNA-seq data from human epidermis. Each point represents a single cell and color coding of cell type populations are shown adjacent (Figure 3B). B. Bar plots showing the fraction of each cell type from the scRNA-seq data from human epidermis. Color-coding of cell types correspond to the colors of the single cells in Figure 3A. C. UMAP plot of clustered scRNA-seq data from mouse skin. Each point represents a single cell and color coding of cell type populations are shown adjacent in Figure 3D. D. Bar plots showing the fraction of each cell type from the scRNA-seq data from mouse skin. Color-coding of cell types correspond to the colors of the single cells in Figure 3C. E. UMAP plot of integrated scRNA-seq data from human epidermis and mouse skin. Each point represents a single cell and color coding of cells indicates the species the cells were obtained from (human = green; mouse = purple). F. UMAP plot of integrated scRNA-seq data from human epidermis and mouse skin. Each point represents a single cell and color coding of cell type populations and collapsed superpopulations are shown in the adjacent legend. G, H. Bar plots showing the fraction of cell types estimated in GTEx skin RNA-seq samples from human epidermis scRNA-seq (G) and mouse skin scRNA-seq (H). I. Heatmap showing the correlation of GTEx skin cell population estimates from mouse skin scRNA-seq at high and collapsed resolutions (rows) and human skin (columns). Color coding of heatmap scales from red, indicating negative and low correlation in estimates, to blue, indicating positive and high correlation in estimates. Significance is indicated with asterisks. J, K. Scatter plots of estimated cell compositions across 860 GTEx skin samples deconvoluted using human scRNA-seq for human keratinocyte 14 population versus mouse stem cell of epidermis population (J) and keratinocyte 1, 5, 14, 711 population versus collapsed mouse epidermal cell populations (K).

We next compared the cellular composition estimates of 860 GTEx bulk skin RNA-seq samples^1^ obtained by deconvolution using human gene expression signatures to mouse gene expression signatures (Figure 1A, “proof-of-concept”). We obtained signature genes for each cell type identified in scRNA-seq from human skin (i.e. signature genes from each of 9 dermis cell types) and mouse skin (i.e. signature genes from each of 6 skin cell types) (Table S3) and used CIBERSORT to deconvolute the 860 GTEx skin RNA-seqs (Figure 3G,H). Given the presence of the three superpopulation clusters observed in the mouse and human scRNA-seq integration analysis (Figure 3F), similar to liver, we investigated how resolution impacted the correlation between human and mouse signature gene estimates. We independently collapsed the high resolution human epidermis (9 cell types) and the high resolution mouse skin (6 cell types), by summing the estimates across the cell types in each of the three distinct superpopulations (Table S4). We then calculated all pairwise-correlations between each of the estimated cell populations in the 860 GTEx skin samples from human estimates (high and collapsed) with the estimated cell populations from mouse (high and collapsed resolution) (Figure 3I, Figure S3). Using the integration analysis (Figure 3F) as a guide, we examined the similarity of estimates from human and mouse cell populations mapping to each of the three superpopulations. First, we examined the similarity of human cell types in Superpopulation 1 (Keratinocyte 14, Keratinocyte 5, Keratinocyte 711, Keratinocyte 1, cornified envelope, and collapsed estimates of these cell types) and mouse cell types in Superpopulation 1 (epidermal cell, basal cell, stem cell of epidermis, outer bulge, and collapsed estimates of these cell types) (Figure 3F; dark purple shading). We observed the human keratinocyte population 14 had a strong positive correlation with the mouse stem cell of the epidermis estimates (R = 0.89; p = 2.4 x 10^-103^) (Figure 3I,J). We also found that collapsed mouse epidermal cell estimates were correlated with collapsed human keratinocyte population estimates (R = 0.44, p = 1.19 x 10^-43^) (Figure 3I,K). These results indicate that despite differences in annotations, estimates from mouse and human cell types mapping to the epidermal cell superpopulation are highly correlated. Second, we examined the similarity of human cell types in superpopulation 2 (fibroblast 0, fibroblast 3, fibroblast 4, and collapsed estimates of these cell types) and the single mouse cell type (inner bulge) in this cluster (Figure 3F; light purple shading). We found that human fibroblast (high resolution and collapsed) estimates were not correlated with the mouse inner bulge cell population estimates (Figure 3I), indicating that, despite similar enough global gene expression patterns for the human fibroblasts and mouse inner bulge cells to cluster together, their signature genes distinguish them as different cell types during deconvolution. Third, we examined the similarity of the human cell type (lymphocyte) and mouse cell type (leukocyte) in superpopulation 3 (Figure 3F; pink shading). Similar to the liver estimates, mouse and human leukocyte estimates were not correlated (Figure 3I), likely due to known species differences in immune cells. As we observed in liver, we confirmed that technical and biological differences influence cell estimate performance, however overall cell composition estimates derived from human and mouse skin signature genes are correlated, supporting our ability to use mouse scRNA-seq as an alternative to human scRNA-seq for the deconvolution of GTEx tissues.

### Cellular deconvolution of GTEx tissues reveals surprising levels of heterogeneity

To understand the extent to which the mouse signature genes obtained from cell types across 14 tissues were able to distinguish between the 28 GTEx tissues, we extracted the expression of the signature genes (Table S1,3) across the 6,829 bulk GTEx RNA-seq samples and visualized how the samples clustered (Figure 4A). We observed that the mouse signature genes were able to differentiate between the human GTEx organs, as well as illustrated the existence of organ substructures delineating heterogeneity in tissues belonging to the same organ. For example, tissues from the same organ clustered closely together and distinctly from other organs, including the heart tissues (atrium and ventricle), brain tissues (cortex, frontal cortex, hippocampus, anterior cingulate cortex, amygdala, substantia nigra, spinal cord, putamen, nucleus accumbens, caudate, and hypothalamus), adipose (visceral and subcutaneous), and colon (sigmoid and transverse). Of note, within the brain we also observe clustering according to zonation, including clustering of samples from the cerebellum and cerebellar hemisphere, as well as clustering of samples from the frontal cortex, cortex, and anterior cingulate cortex. These results suggest that signature gene capture both organ differences, as well differences that exist in tissues from the same organ that hint at tissue substructures driven by sample heterogeneity.

**Figure 4:**
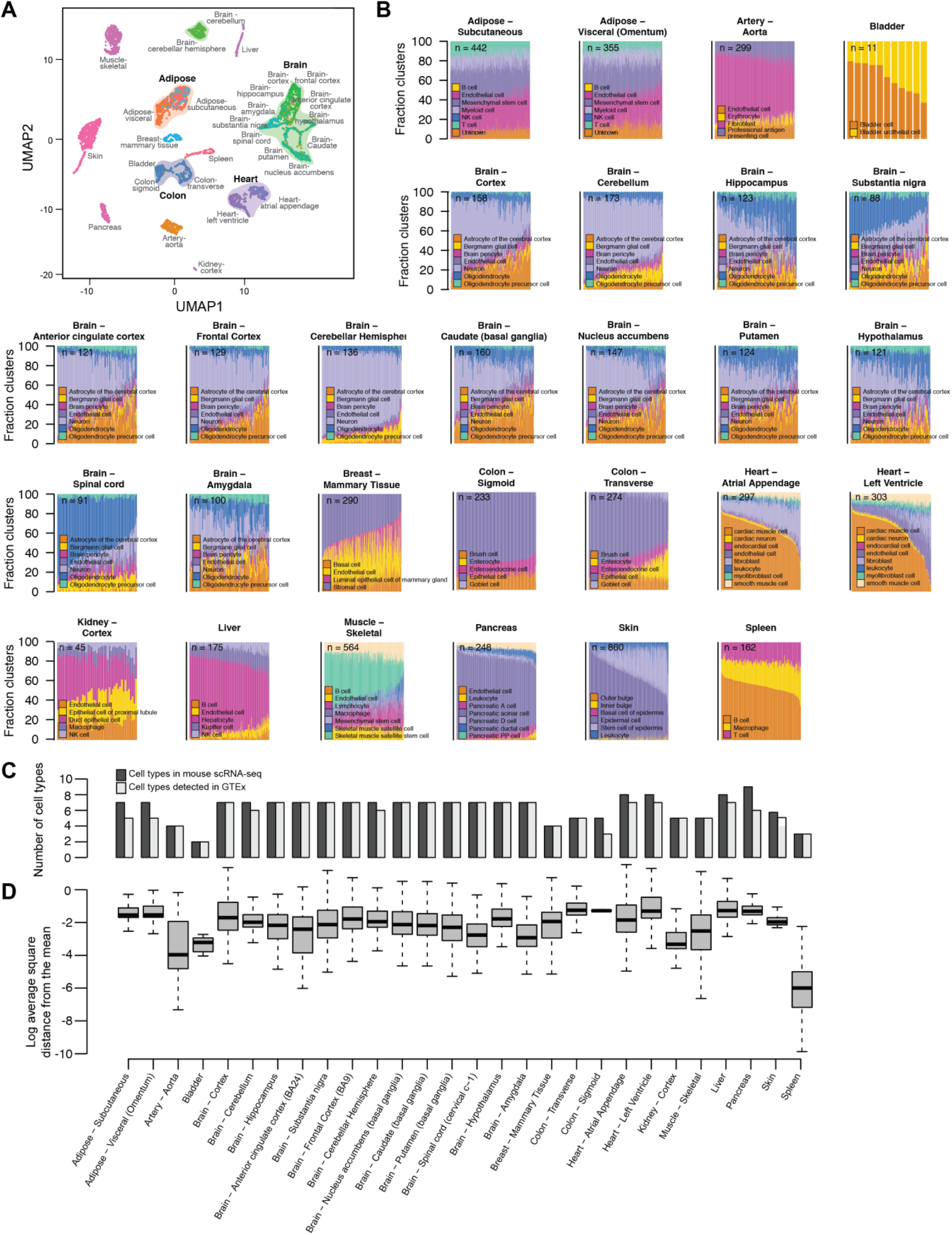
Cellular deconvolution of 28 GTEx tissues. A. UMAP using the expression of all scRNA-seq derived signature genes across the 28 GTEx tissues B. Stacked bar plots showing the fraction of cell types estimated in GTEx RNA-seq samples from mouse scRNA-seq. A colorblind-friendly version of this figure is shown in Figure S4. C. Bar plots comparing the number of cell types discovered in mouse scRNA-seq (light grey) vs. the number of these cell types that were estimable for each GTEx tissue D. Box plots showing per RNA-seq sample the distribution of the log2 average square distance from the mean estimated cellular compositions for each GTEx tissue

To understand the cellular heterogeneity of 28 GTEx tissues (Figure 1A, “cellular deconvolution”), we used the signature genes from 14 mouse tissue types (Table S1,3) to perform cellular deconvolution of 28 GTEx tissues from 14 organs (Figure 4B; Table S1, S5-20), where the number of samples for each GTEx tissue varied from 11 (bladder) to 860 (skin). We found that all samples were well-deconvoluted (P-value < 0.001; CIBERSORT, 1,000 permutations) and that each deconvoluted GTEx tissue contained a variable number of cell types ranging from two (bladder) to seven (brain and heart) (Figure 4C). In ∼30% of the tissues (9 out of 28), we found that not all mouse cell types were estimated, possibly due to the GTEx tissues having been isolated for bulk RNA-seq from a different spatial location than mouse or species differences in cell types. Additionally, the relative distribution of the estimated cell types varied between different samples of the same tissue (Figure 4D). Tissues with the least heterogeneous cell population distributions between samples were aorta and spleen (Table S5,19), whereas those with the most heterogeneous cell population distributions between samples were brain (13 tissues), colon, and left ventricle (Table S8,9,20). Examining the tissues corresponding to the same organ, we noted that some had the same cell types estimated at similar distributions (adipose subcutaneous and visceral), some had the same cell types present at variable proportions (heart atrial appendage and left ventricle; 13 brain tissues), and others had variable cell types present/absent (colon transverse and sigmoid). These results reveal a striking heterogeneity in GTEx tissues that has not been previously appreciated and may be contributing noise to eQTL analyses.

### eQTL analyses using deconvoluted tissues increases power

Since we observed heterogeneity in the relative distributions of cell populations across GTEx RNA-seq samples, we hypothesized that considering the cell population distributions of each sample would improve eQTL analysis by increasing our power to detect novel tissue and/or cell type associations (Figure 1A). We identified 19,621 expressed genes in GTEx liver samples and performed one eQTL analysis not considering cellular heterogeneity (i.e. bulk resolution; Table S21), and three eQTL analysis using cell population estimates as covariates to adjust for cellular heterogeneity (Tables S22-24): 1) considering high resolution human liver estimates (15 cell types; Table S12, 22; Figure 2A); 2) considering collapsed resolution human liver estimates (7 cell types; Table S2,12,23; Figure 2C); and 3) considering low resolution mouse liver estimates (5 cell types; Table S13,24; Figure 2B). Using cell population estimates as covariates we detected many more genes with significant eQTLs (eGenes) than at bulk resolution (Figure 5A). We found that considering high resolution estimates identified the most eGenes (10,117) with 1.3 fold and 3.1 fold more than collapsed and low resolution estimates, respectively. These findings show that conducting eQTL analyses using highly resolved cell population estimates as a covariate significantly increases the power to identify eGenes.

**Figure 5:**
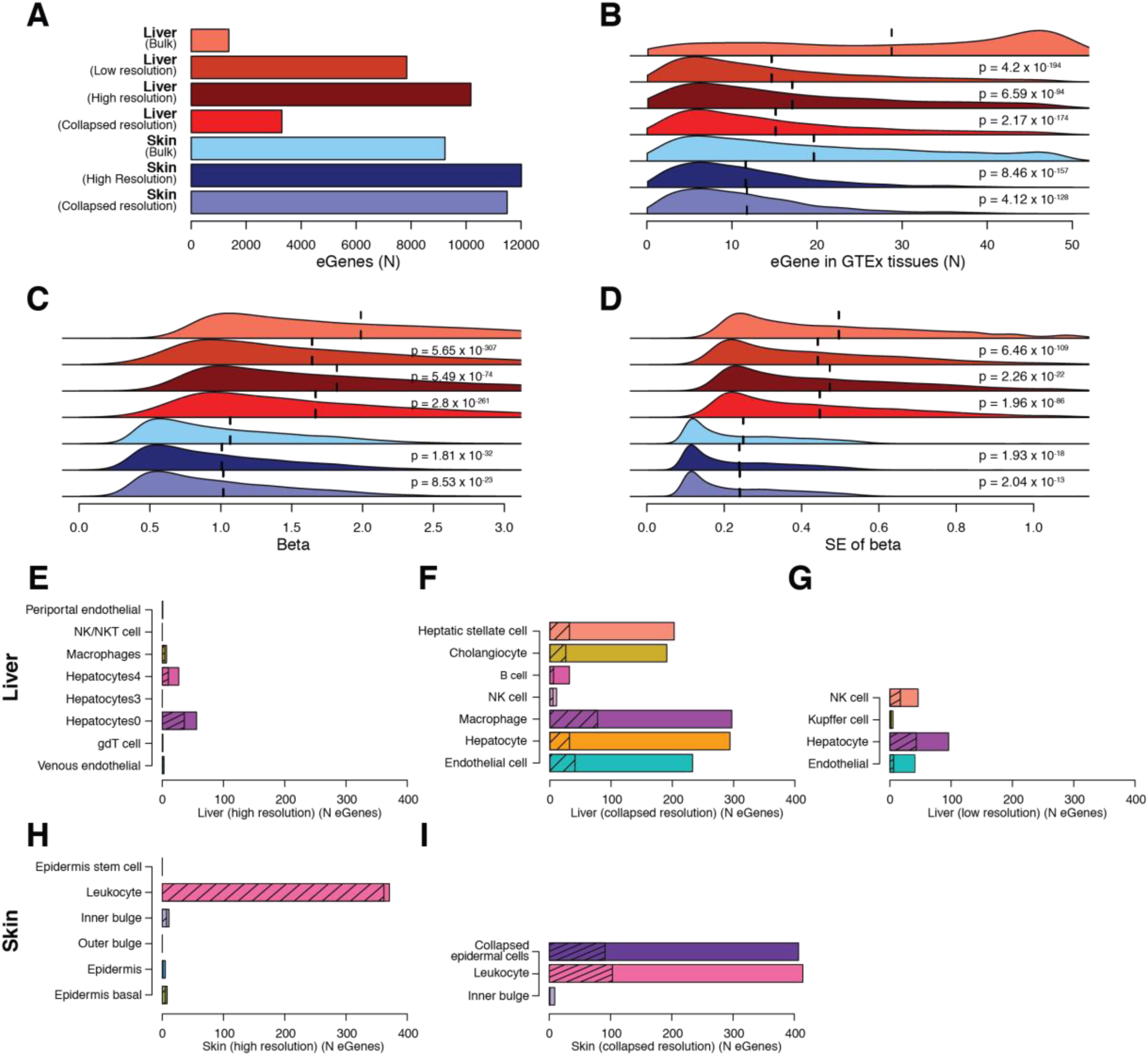
Using cellular deconvolution to discover cell-type-associated eQTLs. A. Bar plot showing the number of eGenes detected in each eQTL analysis from liver (shades of red) and skin (shades of blue). B, C, D. Distributions of (B) number of GTEx tissues where each eGene has significant eQTLs, (C) effect size β and (D) standard error of β in liver and skin. Colors are as in panel A. Vertical dashed lines represent mean values. P-values were calculated in comparison with the bulk resolution analysis for each tissue using Mann-Whitney U test. E-I. Bar plots showing the number of eGenes significantly associated with each cell population considering cell estimates for: liver high resolution (E), liver collapsed resolution (F), liver low resolution (G), skin high resolution (H), and skin collapsed resolution (I). Total number of eGenes for each cell type indicates the cell type is significantly associated and the hashed number of eGenes for each cell type indicates the association is cell-type-specific (e.g. only significant in that cell type). In cases where a given cell type had no significant association, the bar is not shown.

Given the differences in the number of detected eGenes based on cell-type resolution, we hypothesized that eGenes detected at low powered resolutions (bulk and collapsed resolution) commonly shared eQTLs with other GTEx tissues (i.e. tissue-neutral) and the eGenes detected using higher powered resolutions had more tissue-associated eQTLs (i.e. less frequently in other GTEx tissues). For each resolution, we calculated the number of GTEx tissues in which each eGene has eQTLs. We observed that eGenes identified using cell populations as covariates in general were more tissue-associated than eGenes detected at bulk resolution. Compared to bulk resolution, high resolution eGenes were the most tissue-associated (p = 4.2×10^-194^; Mann-Whitney U test), then low resolution eGenes (p = 2.17×10^-174^; Mann-Whitney U test), and collapsed resolution was the least tissue-specific (p = 6.59×10^-94^; Mann-Whitney U test) (Figure 5B), showing that the resolution of cell population estimates used as covariates is correlated with the power of the study to identify tissue-associated eGenes.

Furthermore, using cell populations as covariates resulted in decreased effect size (β) (Figure 5C) and standard error (SE) of β (Figure 5D), where relative to bulk resolution, the higher the resolution of the eQTLs, the smaller the β and SE of β. However, in general the β values for the top hit for each gene were highly correlated between eQTLs detected using cell populations and eQTLs detected without using cell populations (r > 0.975, Figure S6A-C). By performing a permutation test, we excluded that the detection of a larger number of eQTLs using cell populations was due to the fact that a larger number of covariates was used (Figure S7). These results indicate that using cell population distributions as covariates overall reduces the noise, thereby potentially increasing our power to identify eQTLs.

### Resolution of deconvoluted tissues impacts the number of identified cell-type-associated regulatory variants

To examine if some of the eQTLs identified using cell population estimates as covariates were cell-type-associated, we used a statistical interaction test^16–18^ to assess if modeling the contribution of a cell type significantly improved the observed association between genotype and gene expression. Interaction tests were performed on all independent pairs of eGenes and corresponding lead eQTLs using liver cell type estimates from the high, collapsed, and low resolution as interaction terms. Overall, across the high, low, and collapsed resolutions we respectively detected 74, 121, and 528 cell-type-associated eGenes (i.e., eGene is more significant considering cell type estimates but associated with one or more cell type(s); FDR-corrected p-values < 0.1, χ^2^ test, Figure 5E-G) and 54, 68, and 220 cell-type-specific eGenes (i.e. eGene is associated with only one cell type; Figure 5E-G). We investigated if relative cell abundance influenced our ability to detect cell-type-associated eGenes (i.e., is there more power for high abundance cells) and determined that it did not play a factor (Figure S8). Further, we noted by using low resolution and collapsed resolution cell populations, we respectively detected 1.6 and 7.1 times more cell-type-associated eQTLs than high resolution cell populations (respectively, p = 1.9 x 10^-7^ and 7.3 x 10^-250^, Fisher’s exact test, Figure 5E-G). While initially counter-intuitive to the previous evidence showing higher resolution eGenes are more tissue-specific (Figure 5B) and have decreased noise (Figure 5C,D), it is possible we identify a greater number of cell-type-associated eGenes using low resolution cell population estimates due to prevention of the dilution of eQTL signals between shared cell types, as might occur in cases where a regulatory variant has similar effects across similar cell types. Overall, these results suggest that accounting for cellular heterogeneity between samples allows for the discovery of novel cell-type-associated (and cell-type-specific) eQTLs.

### eQTL analysis of deconvoluted GTEx skin confirms ability to identify cell-type-associated regulatory variants

To further investigate the impact of using cell populations on power to identify novel eGenes and cell-type-associated eQTLs, we conducted eQTL analyses using the GTEx tissue (skin), which includes the largest number of RNA-seq samples (Figure 4B). Although we deconvoluted 860 skin RNA-seqs using signature genes from high resolution mouse skin scRNA-seq (6 cell types; Figure 3B), only 749 had corresponding genotypes from 510 distinct individuals. We identified 24,029 expressed genes in the 749 skin RNA-seq samples with corresponding genotypes and performed three eQTL analyses: 1) without considering cell population distributions (bulk resolution) (Table S25); 2) considering high resolution mouse skin cell estimates (6 cell types; Table S17, 26; Figure 3C); and 3) considering collapsed resolution mouse skin cell estimates (3 cell types; Table S17,4,27; Figure 3C,F). Using cell (high and collapsed) population distributions as covariates, respectively, we detected a 30% and 24% increase in eGenes with significant eQTLs (12,011 and 11,497 compared with 9,232, Figure 5A). Similar to our observation in liver, we found that eGenes specific for the eQTL analysis performed using high and collapsed cell populations as covariates, respectively, had eQTLs in fewer tissues than eGenes detected at bulk resolution (p = 8.46 x 10^-157^; p = 4.12 x 10^-128^, Mann Whitney U test; Figure 5B), had a decreased effect size β (p = 1.81 x 10^-32^; p = 8.53 x 10^-23^, Mann Whitney U test, Figure 5C), and had decreased standard error (SE) of β (p = 1.93 x 10^-18^; p = 2.04 x 10^-13^, Mann Whitney U test, Mann Whitney U test; Figure 5D). We also observed that the β values for the top hit for each eGene were highly correlated between eQTLs detected using high and collapsed cell populations and eQTLs detected without using cell populations (r = 0.994; r = 0.996, Figure S6D). Further, at high resolution we detected 384 cell-type-associated eGenes (FDR-corrected p-values < 0.1, χ^2^ test, Figure 4H) and 375 cell-type-specific eGenes (FDR-corrected p-values < 0.1, χ^2^ test, Figure 5H), which were predominantly associated with leukocytes, while at collapsed resolution we detected 511 cell-type-associated eGenes (FDR-corrected p-values < 0.1, χ^2^ test, Figure 5I) and 220 cell-type-specific eGenes (FDR-corrected p-values < 0.1, χ^2^ test, Figure 4I), associated with both the collapsed epidermal cell population and leukocytes (Superpopulations 1 and 3; Figure 3F). We hypothesize that substantially fewer cell-type-specific associations were observed in the high resolution epidermal cell types (epidermal cell, basal cell, stem cell of epidermis, outer bulge; Figure 5H) compared with the collapsed epidermal cells (Figure 5I), because of a dilution of signal between similar cell types. The relatively large number of cell-type-associated eGenes in skin compared with the liver could be reflective of sample size differences between the two tissue (749 and 153, respectively) impacting power to detect eGenes. These results show that even in eQTL studies using large sample sizes, accounting for cellular heterogeneity results in the detection of thousands more eGenes, which tend to show cell-type-associated differential regulation.

### Colocalization identifies cell-type-associated regulatory variants are associated with specific skin diseases

To explore the functional impact of the cell-type-associated eQTLs identified in skin, we examined their overlap with GWAS signals for skin traits and disease. From the UK Biobank, we extracted GWAS summary statistics for 23 skin traits where the cell types identified from skin scRNA-seq (Figure 6A) likely played a role in the traits (Table 28) and grouped them into seven categories based on trait similarity: 1) malignant neoplasms, 2) melanomas, 3) infections, 4) ulcers, 5) congenital defects, 6) cancer (broad definition, non-malignant neoplasm), and 7) unspecified skin conditions. As the three collapsed skin superpopulations identified the most cell-type-associated eGenes, we performed colocalization of the eQTLs identified using the collapsed resolution cell estimates (Table S4) and skin GWAS loci to identify shared causal variants using coloc^19^ and examining instances with PP4 > 0.5 (PP4, posterior probability of the colocalization model having one shared causal variant). We identified 394 variants that showed evidence of colocalization (Table 28). These results show that we could identify hundreds of skin eQTLs that likely share a causal variant with skin GWAS traits.

**Figure 6:**
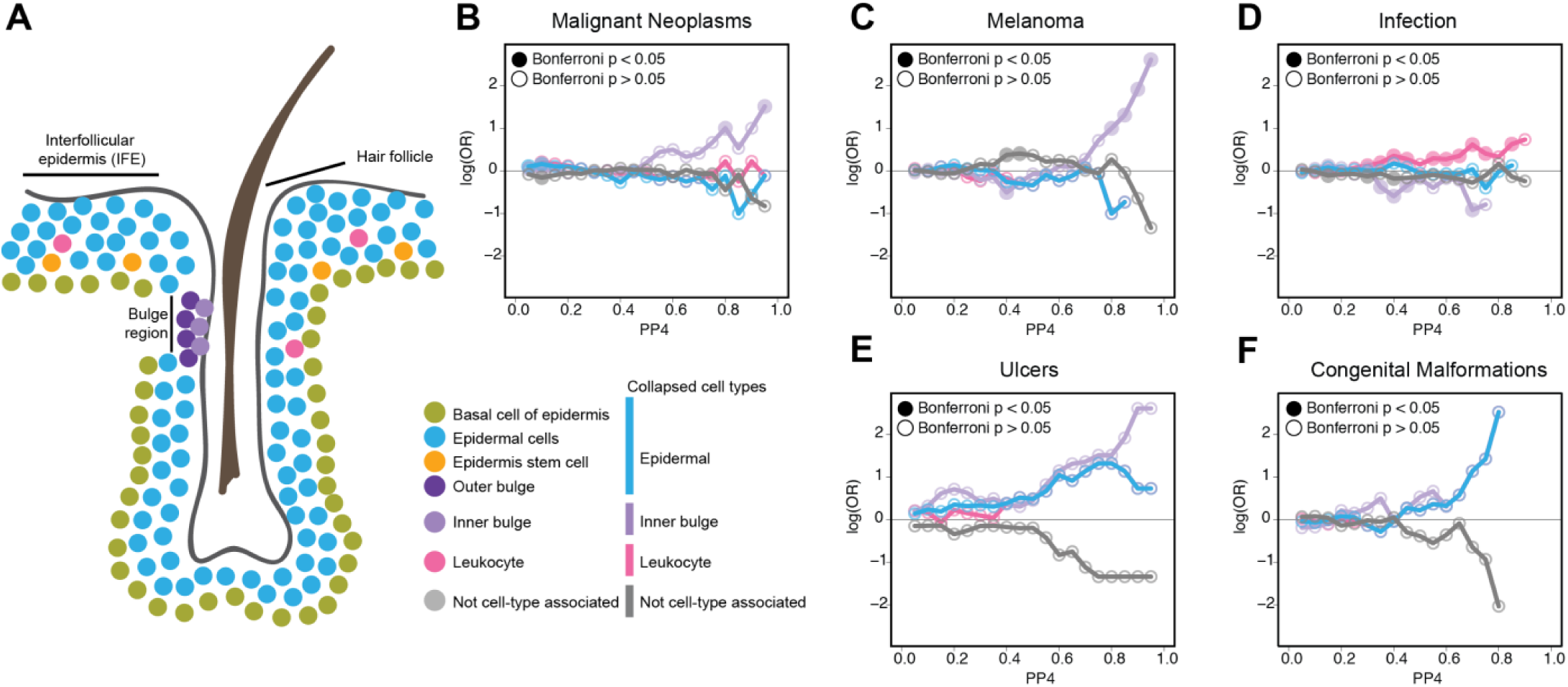
Colocalization of cell-type-associated skin eQTLs with skin GWAS traits. A. Cartoon describing the approximate organization of cell types identified in scRNA-seq from skin. Colors used for each cell type are used throughout Figure and described in the adjacent legend. B-F. Line plots showing the enrichment of cell-type-associated eQTLs in various GWAS traits: malignant neoplasms (c), melanoma (D), infection (e), ulcers (f), and congenital malformations (g). Enrichment is plotted as the log(OR) (y-axis) over all probabilities of the eQTL signal overlapping (0 = not overlapping – 1 = completely overlapping) with the GWAS signal (x-axis). Lines are colored following color coding of each cell type from 5A.

We next asked if skin GWAS traits were enriched for eQTLs that are associated with distinct cell types. We tested the enrichment of cell-type-associated eQTLs at multiple PP4 thresholds and found malignant neoplasms and melanomas were enriched for eQTLs associated with keratinocyte stem cells from the inner bulge (p = 1.13 x 10^-3^, p = 2.82 x 10^-4^ Fisher’s Test; Figure 6B,C), and infections were enriched for eQTLs associated with leukocytes (p = 9.69 x 10^-3^ Fisher’s Test; Figure 6D). We did not observe a significant enrichment of cell-type-associated eQTLs in ulcers (Figure 6E), congenital malformations (Figure 6F), cancer (broad definition), or unspecified skin conditions. It is unclear if this is to be expected, as it is possible other cell types not estimated may be contributing to the diseases or in the case of congenital malformations, it is possible that expression differences impacting congenital malformations may be functioning during development and not detectable in adult skin. Overall, these results suggest that GWAS lead variants are commonly cell-type-associated regulatory variants, indicating that onset or progression of human disease and traits may be controlled at the cell type level.

We next sought to specifically examine the eGenes that most strongly colocalized with malignant neoplasms or melanoma (PP4 ≥ 0.8), as bulge stem cells have been implicated in playing a role in cancer^20–25^. We found six eGenes not previously associated with skin cancers with eQTLs significantly associated with inner bulge stem cells, including: 1) *BRIX1*, which has been found to play a role in cancer progression^26^; 2) *RP11-875011.*1, an antisense gene, which has not previously been implicated in cancer, however antisense genes are thought to contribute to the regulation of human cancers^27^; 3) *MUL1*, which has been associated with the progression of human head and neck cancer^28^; 4) *PMS2P3*, has been implicated in affecting survival in pancreatic cancer^29^; 5) *FTH1*, which has been shown to be involved in regulating tumorigenesis^30, 31^ and whose increased expression in keratinocytes may be in response to stress^32, 33^; and 6) *CNTN2,* which is involved in cell adhesion and has been implicated in tumor development^34, 35^. The identification of these disease-associated eGenes supports our ability to identify cell-type-associated eQTLs whose functions are congruent with playing a role in the etiology of cancer. Together these results show that conducting eQTL studies accounting for cellular heterogeneity can identify the likely causal cell-type-associated variants and genes underlying GWAS disease loci.

## Discussion

Human scRNA-seq data representative of all tissues in GTEx that could be used to deconvolute the more than 10,000 GTEx bulk RNA-seq samples does not yet exist. As the Tabula Muris resource of mouse scRNA-seq from 20 organs was recently released^11^, we sought to determine if mouse signature genes obtained from scRNA-seq could be used as an alternative for human signature genes for cellular deconvolution of GTEx RNA-seq samples. Using scRNA-seq from both mouse and human for two proof-of-concept tissues (liver and skin), we derived signature genes and used these expression profiles to deconvolute GTEx liver and skin RNA-seq samples. In general, human and mouse estimates between the two proof-of-concept tissues were comparable, where discrepancies in cell composition estimates between the two species primarily resulted from technical and subtle immunological differences. Specifically, in both liver and skin, technical differences impact the resolution at which cellular composition can be estimated, including: 1) the number of cells captured and subjected to scRNA-seq; and 2) tissue sampling methodology. Further, differences in cell composition estimates for immune cells were observed most likely due to immunological differences between the two species. These differences highlight that high resolution scRNA-seq (more cells/cell types sampled from diverse zones) is key to identifying and estimating the composition of highly specialized and rare cell types. For these reasons, the cell composition estimates we obtained from CIBERSORT using mouse-derived signature genes from proof-of-concept liver and skin scRNA-seq may still be missing cell types not captured in the Tabula Muris resource. An additional challenge we found that influenced our ability to compare cell composition estimates was the scRNA-seq cell annotations in human and mouse skin did not use consistent naming conventions, thus it was not immediately clear how to compare cell estimates across the studies. We were able to overcome this challenge by integrating the mouse and human scRNA-seq, which allowed us to infer three similar superpopulations of cell types across the two species based on gene expression.

To examine cellular heterogeneity across the GTEx resource, we used the signature genes obtained using scRNA-seq from 14 mouse tissue types to deconvolute 6,829 GTEx RNA-seq samples mapping to 28 tissues from 14 organs. We found that GTEx tissues exhibit substantial cellular heterogeneity, with the number of cell types ranging from two in bladder to seven in brain and heart. Additionally, some of the tissues, including brain, colon, and left ventricle, showed highly variable proportions of estimated cell types between samples, contributing to intra-tissue cellular heterogeneity. Together, these results reveal a source heterogeneity in GTEx tissues that has not been previously considered and may contribute to reduced power to detect eQTLs.

While genetic association studies performed by GTEx have identified a wealth of novel insights into how human genetics function across bulk tissues^1^, these analyses have not considered how cellular heterogeneity can confound these studies through biasing or even masking cell-type-specific signals. We found that considering cellular heterogeneity significantly improved eQTL analyses by increasing power to detect lower effect size genetic associations, as well as by identifying cell-type-specific associations that were masked in analyses using bulk RNA-seq data from the same samples. Further, we found resolution of cell heterogeneity influenced eQTL results, where considering high resolution estimates identified substantially more eQTLs than using lower resolutions (low resolution or collapsed resolution); however, high resolution cell estimates identified fewer cell-type-associated genetic associations than lower resolutions. It is possible this decrease in associations may be due to a dilution of signal between similar cell types. Our observations suggest these two resolutions should both be used to power eQTL analyses in complementary ways: 1) high resolution estimates to power association analyses to discover lower effect size eQTLs; and 2) collapsed resolution estimates to identify cell-type associated eQTLs. We further show that cell-type-associated eQTLs colocalize with lead variants from relevant GWAS traits, highlighting a potential path forward for understanding the impact of genetic variation on mechanisms underlying complex traits.

Overall, we demonstrate that while efforts to generate a resource of scRNA-seq data from human tissues^36^ are in progress, QTL studies using human bulk RNA-seq data could utilize readily available mouse-derived signature genes to estimate cellular heterogeneity and optimize power to identify cell type-specific genetic associations. As the Tabula Muris resource does not represent all of the human GTEx tissues (28 of 53) it is possible that scRNA-seq resources from other mammalian species could be used to deconvolute the non-represented GTEx tissues. Our study further emphasizes that the straightforward approach of taking tissue heterogeneity into account when conducting genetic association studies has the potential to greatly expand our understanding of the functional impact of genetic variation on molecular and complex human traits.

## Methods

### Mouse single cell transcriptome profiles from 14 mouse organs from Tabula Muris

Single cell transcriptome profiles from 14 mouse organs were used in this study^11^. Briefly, transcriptome profiles were generated from three female and four male mice (C57BL/6JN; 10-15 month-old) from: aorta, atrium, bladder, brain nonmicroglia, colon, fat, kidney, liver, mammary gland, muscle, pancreas, skin, spleen, ventricle (Table S1). Upon extraction of these organs from the mice, single cell transcriptomes were generated by first sorting by fluorescence-activated cell sorting (FACS) for specific populations (FACS method; SMART-Seq2 RNAseq libraries). We downloaded the normalized gene expression and annotated single-cell clusters from each organ as Seruat^12^ R objects (https://figshare.com/articles/Robject_files_for_tissues_processed_by_Seurat/5821263/1).

### Processing of scRNA-seq from human liver

10X Genomics formatted BAM files from five human total liver homogenate samples^6^ were downloaded (GEO accession: GSE11546) and converted to fastq files using 10X bamtofastq (https://support.10xgenomics.com/docs/bamtofastq). Converted fastq files were then processed using cellranger count utility to generate gene expression count matrices, then the five processed liver samples were merged using cellranger aggr utility.

### Annotation of the cell populations present in human liver scRNA-seq data

Analysis of scRNA-seq from human liver^6^ were conducted following the same approach used to annotate mouse organs^11^. Cells with fewer than 500 detected genes or cells with fewer than 1,000 UMI were filtered from the data, resulting in 8,119 cells analyzed from human liver. Gene expression was then log normalized and variable genes were identified using a threshold of 0.5 for the standardized log dispersion. Principal component analysis (PCA) was performed on the variable genes and significant PCs. Clustering was performed using a shared-nearest-neighbor graph of the significant PCs and single cells were visualized using Uniform Manifold Approximation and Projection (UMAP). Cell populations were then annotated based on the expression of known liver marker genes^11^.

### Collapsing liver cell population estimates

To collapse similar cell populations in GTEx liver samples, we examined the UMAP from high resolution human liver scRNA-seq (Figure 1B) and compared to the UMAP from low resolution mouse liver scRNA-seq (Figure 1C) to identify broader/lower resolution classifications of cell types present in the liver (Table S2). We identified populations in the human liver scRNA-seq that were similar (e.g. Hepatocyte populations 0, 1, 3, and 4; Figure 1B) with a corresponding population in the mouse liver scRNA-seq (e.g. Hepatocyte; Figure 1C). For populations identified in human not present in mouse, we did not perform any collapsing.

### Annotation of the cell populations present in skin scRNA-seq data

*Human*: Skin scRNA-seq^12^ gene expression data and cell annotations for 8,388 cells (Figure S2A) were downloaded from http://dom.pitt.edu/rheum/centers-institutes/scleroderma/systemicsclerosiscenter/database/. Cells with fewer than 200 detected genes were filtered from the data. Gene expression was then log normalized and variable genes were identified using a threshold of 0.5 for the standardized log dispersion. Principal component analysis (PCA) was performed on the variable genes and significant PCs. Clustering was performed using a shared-nearest-neighbor graph of the significant PCs and single cells were visualized using Uniform Manifold Approximation and Projection (UMAP). The single cells were then annotated using provided cell annotations and validated using marker gene expression. As the human skin scRNA-seq contained cell types belonging to various layers of the skin, whereas the mouse scRNA-seq was enriched for epidermal cells, we extracted only the 5,670 human cells belonging to the epidermal layer of the skin. We then reanalyzed the subsetted data following the above methods by performing PCA, reclustering, and visualization using UMAP. *Mouse*: Tabula Muris cell annotations (Figure S2D) were confirmed by examining marker gene expression for epidermal cells, basal cells of the epidermis (Krt1High), stem cells of the epidermis (Top2aHigh), leukocytes (Lyz2High), and keratinocyte stem cells (Cd34High). While Tabula Muris annotated a single keratinocyte stem cell population, we reannotated this population by distinguishing between: 1) inner bulge cell population exhibiting Dkk3High and ITGA6Low expression; and 2) outer bulge cell population exhibiting Fgf18High and ITGA6High expression.

### Deconvolution of complex tissues using CIBERSORT

*Identification of signature genes from single cell populations:* For 16 scRNA-seq datasets from human liver, human skin, and scRNA-seq from 14 mouse organs, we obtained gene expression signatures for each annotated cell type (Table S3) and used as input into CIBERSORT_3_ to estimate the cellular composition of GTEx adult tissues (Table S1). For each tissue, we identified differentially expressed genes using Seurat FindMarkers and then extracted the top 200 most significantly overexpressed genes (adjusted p-value < 0.05; average log2 fold change > 0.25) for each of the annotated scRNA-seq cell types (gene expression signatures). For signature genes obtained from mouse scRNA-seq, we converted the mouse genes to their human orthologs using the biomaRt database^37, 38^. The final gene signature sets only included mouse signature genes that also had a human ortholog. For a given signature gene set: 1) if a mouse gene had more than one human ortholog, only one human ortholog was retained in final signature set; and 2) if different mouse genes corresponded to the same human ortholog, only unique human orthologs were retained in the final signature set.

*Cell composition estimation*: The mean expression levels of the signature genes were used as input for CIBERSORT to calculate the relative distribution of the cell populations of 28 GTEx tissues from 14 organs (Table S5-20). CIBERSORT (https://cibersort.stanford.edu/) was run with default parameters using the TPM values for the signature genes identified from scRNA-seq in all RNA-seq samples from the analogous GTEx tissue (https://gtexportal.org/home/datasets) (Table S1). To determine the cell types detected in GTEx compared to the cell types modeled from mouse, we classified a given cell type as estimable in GTEx as those with CIBERSORT estimates greater than 0.05% in more than 5% of RNA-seq samples from a given GTEx tissue. To estimate cellular heterogeneity across GTEx RNA-seq samples, heterogeneity was measured as the average square distance from the mean for each GTEx tissue. We further examined how time from death or withdrawal of life-support until each tissue sample was fixed/frozen (i.e. ischemic time) is associated with cellular heterogeneity and we did not observe a consistent trend between ischemic time and cellular heterogeneity (Figure S**5**). GTEx organs are defined as the regions from which tissues are sampled (variable name SMTS from sample attributes data table; phv00169239.v7.p2) and GTEx tissues are defined by the distinct area of the organ where the tissue was taken (variable name SMTSD from sample attributes data table; phv00169241.v7.p2). For example, samples from the GTEx organ, colon, is comprised of two tissues: sigmoid colon and transverse colon.

### Harmonization of human and mouse scRNA-seq

To harmonize scRNA-seq from human and mouse liver and skin, mouse genes for each tissue scRNA-seq dataset were first converted to their human orthologs using the BioMart database^37, 38^. Mouse and human scRNA-seq were then harmonized by identifying genes that anchor the two datasets using Seurat FindIntegrationAnchors and using these anchors to integrate the datasets using Seurat IntegrateData. Integrated datasets were then visualized using UMAP and corresponding cell types were identified by examining overlap of mouse and human cells.

### Detecting eQTLs using a linear mixed model

To detect eQTLs, we obtained gene TPMs for 153 liver bulk RNA-seq samples and 749 skin bulk RNA-seq samples (sun-exposed and not sun-exposed) from the GTEx V.7 website (https://gtexportal.org/home/) and downloaded WGS VCF files from dbGaP (525 individuals, phs000424.v7.p2). Only genes with TPM > 0.5 in at least 20% samples were considered (19,621 genes in liver and 24,029 in skin). Gene expression data was quantile-normalized independently for each tissue type. For all eQTL analyses, we used the following covariates: age, sex and the first five genotype principal components (PCs) calculated using 90,081 SNPs in linkage equilibrium^39^. Since some subjects had two skin samples (one sun-exposed and one not sun-exposed), we employed a linear mixed model (LMM) for eQTL detection, using subject ID as random effect (1|subject_id). We fitted LMMs using the lme4 package (https://www.jstatsoft.org/article/view/v067i01/0)) to detect eQTLs in skin, described in the following model^18^:

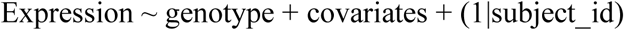

For liver, we used sex as random effect (1|sex) to fit an LMM analogous to the skin eQTL analysis method, described in the following model:

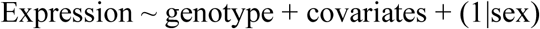

We calculated associations with all variants (minor allele frequency > 1%) ± 1 Mb around each expressed gene. For each gene, we Bonferroni-corrected p-values and retained the lead variant. To detect eGenes, we used Benjamini-Hochberg FDR at 10% level on all lead variants.

### Improved eQTL detection using cell population distributions as covariates

We repeated eQTL detection adding cellular compositions as covariates to the LMMs described above using the following model:

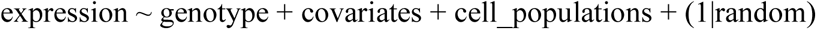

The “cell_populations” term denotes the relative cell population distributions in each tissue and (1|random) is each tissue’s random effect (1|subject_id in skin samples; 1|sex in liver samples, see: Detecting eQTLs using a linear mixed model).

Specifically, we conducted three eQTL analysis for the liver using human high resolution, human collapsed and mouse low resolution cell populations as covariates. Since several cell types were detected at very low frequency, we only used a subset of the cell types described in Figure 4: 1) for human high resolution: periportal sinusoidal endothelial cells, central venous endothelial cells, gdT cells, hepatocytes0, hepatocytes3, hepatocytes4, inflammatory macrophages and NK/NKT cells; 2) for human collapsed resolution: endothelial cells, hepatocytes, macrophages, NK cells, B cells, cholangiocytes, and heptatic stellate cells; and 3) for mouse low resolution: endothelial cells of hepatic sinusoid, hepatocytes, Kupffer cells and NK cells. We conducted two eQTL analysis for skin using mouse high and mouse collapsed resolution cell populations as covariates with the following cell populations: 1) for mouse high resolution: epidermis stem cell, leukocyte, inner bulge, outer bulge, epidermis, and epidermis basal cells; and 2) for mouse collapsed resolution: epidermal cells, leukocyte, and inner bulge cells.

### Detecting cell-type-specific and cell-type-associated eQTLs

In order to detect eQTLs associated with one or more cell types, for each cell population we repeated the eQTL analyses described above (see: Improving eQTL detection using cell population distributions as covariates) by adding an interaction term between the genotype and each cell population (cell*_i_*) estimate to the model (genotype:cell*_i_*). Specifically, for each cell*_i_* estimate, we compared the following two models:

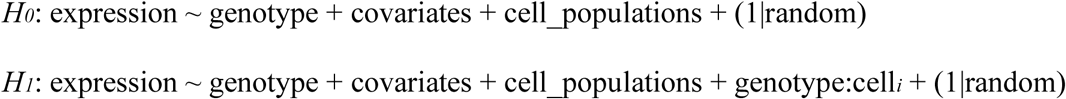

In both models (*H_0_* and *H_1_*), the “cell_populations” term denotes the relative cell population distributions in each tissue and (1|random) is each tissue’s random effect (1|subject_id in skin samples; 1|sex in liver samples, see: Detecting eQTLs using a linear mixed model). In *H_1_*, “genotype:cell*_i_*” is the interaction term between the genotype and cell*_i_* estimate.

To compare the two hypotheses (*H_0_* and *H_1_*), we calculated the difference between the two models using ANOVA and obtained χ2 p-values using the pbkrtest package (https://www.jstatsoft.org/article/view/v059i09).

For each cell*_i_* estimate used as an interaction term in *H_1_*, only eGenes that satisfied two requirements were considered to be associated with cell*i :* a) Benjamini-Hochberg-adjusted χ^2^ p-value <0.1; and b) Δ*_AIC_* = *AIC*_*interaction*_ − *AIC*_*no interaction*_ < 0 (i.e. “genotype:cell*i*” interaction terms that significantly improve the eQTL model). If only one cell population improved the eQTL model, the eQTL was labeled “cell-type specific”; conversely, if more than one cell population improved the eQTL model, the eQTL was labeled “cell type-associated”. We determined the impact of cell population abundance on power to detect cell-type-associated eGenes by examining the distribution of ß, standard error, and P-value for cell-type-associated-eQTLs from each cell population.

### Permutation analysis of liver eQTLs

To test if the detection of more eQTLs using cell populations as covariates was due to improved accuracy of the linear mixed model estimation or was simply associated with an increased number of covariates, for each top hit (defined as the variant with the strongest p-value for each gene), we permuted the cell type distribution across samples, 1,000 times. We obtained the average p-value, beta and standard error of beta across all permutations and compared these values with the measured p-value, beta and standard error of beta for each gene using a paired t-test.

### Colocalization of UK Biobank GWAS for skin traits and eQTLs identified from skin

For each eGene in the skin eQTL analysis deconvoluted using cell type estimates, we extracted the p-values for all variants that were used to perform the eQTL analysis. From the UK BioBank, we obtained summary statistics for 23 skin-related traits (Table S28), where the traits were grouped into seven categories based on shared nomenclature in the trait descriptions: 1) malignant neoplasms; 2) melanoma; 3) infection; 4) ulcers; 5) congenital malformations of the skin; 6) other cancer (non-melanoma or malignant neoplasm); and 7) unspecified. For all the variants genotyped in both GTEx and UK BioBank, we used coloc V. 3.1^19^ to test for colocalization between eQTLs and GWAS signal. For each colocalization test, we considered only the posterior probability of a model with one common causal variant (PP4). Enrichment of the associations was calculated using a Fisher’s Test at multiple PP4 thresholds (0 – 1; by 0.05 bins), where the contingency table consisted of two classifications: 1) *if* the variant was significantly cell-type-associated (FDR < 0.05); and 2) *if* the variant colocalized with the GWAS trait greater than each PP4 threshold.

## Data Availability

Sequence data that support the findings of this study (all Figures) is available for human liver scRNA-seq (GSE11546); for human skin scRNA-seq (http://dom.pitt.edu/rheum/centers-institutes/scleroderma/systemicsclerosiscenter/database/); and for Tabula Muris mouse scRNA-seq (https://figshare.com/articles/Robject_files_for_tissues_processed_by_Seurat/5821263/1). Scripts to process, analyze, and generate Figures from the data is available at https://github.com/mkrdonovan/gtex_deconvolution. The source data underlying all Figures is available in the Source Data file (Tables 1-28).

## Acknowledgements

This work was supported in part by a California Institute for Regenerative Medicine (CIRM) grant GC1R-06673 and NIH grants HG008118, HL107442, DK105541, and DK112155. M.K.R.D. was supported by the National Library of Medicine Training Grant T15LM011271.

## Author information

K.A.F., M.K.R.D., A.D.C., M.D. conceived the study. M.K.R.D and M.D. performed computational analysis. M.K.R.D. performed scRNA-seq data processing and deconvolution analyses. M.D. performed the eQTL analysis. M.K.R.D and M.D. performed colocalization analysis. K.A.F. and A.D.C. oversaw the study. M.K.R.D., M.D., and K.A.F. prepared the manuscript.

