## Supplemental_material for "Cellular deconvolution of GTEx tissues powers eQTL studies to discover thousands of novel disease and cell-type associated regulatory variants"

**SUPPLEMENTAL FIGURES**

**Figure S1. Similarity of eQTL effect size between various resolutions of deconvolution from liver and skin**

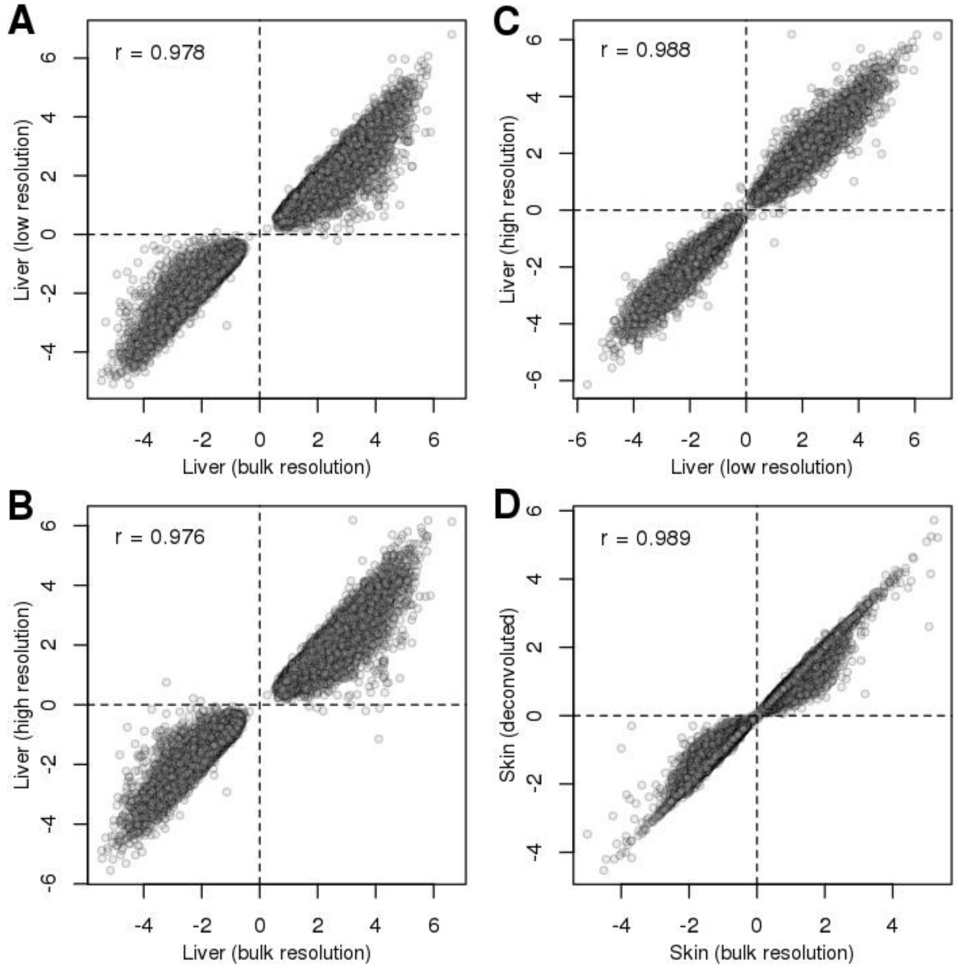


(a) Scatterplots showing the correlation of effect sizes β between different eQTL analyses. For each comparison, the correlation is shown.

### TABLE LEGENDS

**Table S1. Mapping of Tabula Muris mouse scRNA-seq tissues/organs used to deconvolute human GTEx tissues**

GTEx_tissue indicates the GTEx tissue name (Column A) that was deconvoluted using the expression of signature genes from cell populations identified in scRNA-seq from the corresponding mouse organ, indicated in the Tabula_muris_organ column (Column B).

**Tables S2. Expression of signature genes from cell populations identified in 14 scRNA-seq samples**

For each scRNA-seq dataset (14 organs from mouse and one tissue from human), we extracted signature genes from cell populations described in the scRNA-seq. For each organ/tissue (Column A) and species (Column B), the signature gene ensemble IDs (signature_gene_ids; Colum C) are given.

**Tables S3. Scheme used to collapse similar high-resolution liver cell types**

For each of the 15 cell types identified in human liver scRNA-seq, High_resolution_cell_types (Column A) indicates the similar cell types that were collapsed and the collapsed naming scheme is indicated by Collapsed_cell_type (Column B).

**Table S4-18. Cellular composition estimates of bulk RNA-seq from GTEx tissues**

For Tables S4-18, each table represents a different deconvoluted GTEx tissue (15 tables using signature genes from 14 mouse tissue types scRNA-seqs and one table using signature genes from human liver scRNA-seq; Table S2). Each of these tables contain Input.Sample (Column A), which indicates the GTEx bulk RNA-seq sample that was deconvoluted from the tissue captured by each table. In each of these tables, subsequent columns include the proportion of estimated cell populations estimated using signature genes (Tables S2) from corresponding human or mouse scRNA-seq sample (Table S1). In Tables S4-18, each deconvoluted GTEx tissue, GTEx tissues contained within the table, and the tissue type/species signature genes set (Table S2) used for estimation are summarized below:

| Table | Deconvoluted GTEx organ | GTEx tissues included in table | Signature genes |
|---|---|---|---|
| 4 | Artery | Aorta | Aorta/mouse |
| 5 | Heart | Atrial appendage | Heart (subsetted to atrium cells)/mouse |
| 6 | Bladder | Bladder | Bladder/mouse |
| 7 | Brain | Amygdala, Anterior cingulate cortex (BA24), Caudate (basal ganglia), Cerebellar Hemisphere, Cerebellum, Cortex, Frontal Cortex (BA9), Hippocampus, Hypothalamus, Nucleus accumbens (basal ganglia), Putamen (basal ganglia), Spinal cord (cervical c-1), Substantia nigra | Brain-nonmicroglia/mouse |
| 8 | Colon | Sigmoid, transverse | Colon/mouse |
| 9 | Adipose | Subcutaneous, Visceral (Omentum) | Fat/mouse |
| 10 | Kidney | Kidney - Cortex | Kidney/mouse |
| 11 | Liver | Liver | Liver/human |
| 12 | Liver | Liver | Liver/mouse |
| 13 | Breast | Mammary | Mammary/mouse |
| 14 | Muscle | Skeletal muscle | Muscle/mouse |
| 15 | Pancreas | Pancreas | Pancreas/mouse |
| 16 | Skin | Not sun exposed (Suprapubic), Sun exposed (Lower leg) | Skin/mouse |
| 17 | Spleen | Spleen | Spleen/mouse |
| 18 | Heart | Left ventricle | Heart (subsetted to ventricle cells )/mouse |

**Table 19-24. eQTL results**

Six eQTL analyses were performed: 1) GTEx liver at bulk resolution (Table 19); 2) GTEx liver at high resolution (Table 20); 3) GTEx liver at collapsed resolution (Table 21); 4) GTEx liver at low resolution (Table 22), 5) GTEx skin at bulk resolution (Table 23), and 6) GTEx skin with cell populations (Table 24). For each of these tables the ensemble gene ID (Column A), the gene name (Column B), the start (Column C) and end (Column D) of the gene, the gene ID (chromosome_start_end_genotype; Column E), variant position (Column F), reference allele (Column G), alternative allele (Column H), the reference SNP cluster ID (Column I), delta AIC and p-value for covariates included in the model, and FDR are included.

**Table 25. Skin eQTL colocalization with skin GWAS traits results**

For each UK Biobank GWAS trait (Column A) that was colocalized with the eQTLs identified using skin populations, we provide a description of the trait (Column B) and how we collapsed results from similar studies (Column C). This table further describes the gene ensemble IDs (Column D), the gene ID (chromosome_start_end_genotype; Column E), and the number of SNPS tested (Column F). Results of the colocalization include the posterior probability of the model not sharing a signal (PP0; Column G), the posterior probability of only the eQTL having a signal (PP1; Column H), posterior probability of only the GWAS having a signal (PP2; Column I), posterior probability of both the GWAS and eQTL having a signal, but the causal variant is different (PP3; Column J), posterior probability of both the GWAS and eQTL having a shared causal variant (PP4; Column K), and the posterior probability of the SNP (Column L). We also include the p-value for each cell type used as covariates included in the model for the eQTL analysis: Epidermal cell (Column M), Keratinocyte stem cell (Column N), Leukocyte (Column O), and Epidermal stem cell (Column P). Shown are only eGenes with PP4 > 0.2.
